# Protein nanoparticle vaccines induce potent neutralizing antibody responses against MERS-CoV

**DOI:** 10.1101/2024.03.13.584735

**Authors:** Cara W. Chao, Kaitlin R. Sprouse, Marcos C. Miranda, Nicholas J. Catanzaro, Miranda L. Hubbard, Amin Addetia, Cameron Stewart, Jack T. Brown, Annie Dosey, Adian Valdez, Rashmi Ravichandran, Grace G. Hendricks, Maggie Ahlrichs, Craig Dobbins, Alexis Hand, Catherine Treichel, Isabelle Willoughby, Alexandra C. Walls, Andrew T. McGuire, Elizabeth M. Leaf, Ralph S. Baric, Alexandra Schäfer, David Veesler, Neil P. King

## Abstract

Middle East respiratory syndrome coronavirus (MERS-CoV) is a zoonotic betacoronavirus that causes severe and often lethal respiratory illness in humans. The MERS-CoV spike (S) protein is the viral fusogen and the target of neutralizing antibodies, and has therefore been the focus of vaccine design efforts. Currently there are no licensed vaccines against MERS-CoV and only a few candidates have advanced to Phase I clinical trials. Here we developed MERS-CoV vaccines utilizing a computationally designed protein nanoparticle platform that has generated safe and immunogenic vaccines against various enveloped viruses, including a licensed vaccine for SARS-CoV-2. Two-component protein nanoparticles displaying MERS-CoV S-derived antigens induced robust neutralizing antibody responses and protected mice against challenge with mouse-adapted MERS-CoV. Electron microscopy polyclonal epitope mapping and serum competition assays revealed the specificities of the dominant antibody responses elicited by immunogens displaying the prefusion-stabilized S-2P trimer, receptor binding domain (RBD), or N-terminal domain (NTD). An RBD nanoparticle vaccine elicited antibodies targeting multiple non-overlapping epitopes in the RBD, whereas anti-NTD antibodies elicited by the S-2P– and NTD-based immunogens converged on a single antigenic site. Our findings demonstrate the potential of two-component nanoparticle vaccine candidates for MERS-CoV and suggest that this platform technology could be broadly applicable to betacoronavirus vaccine development.

## Introduction

The recent SARS-CoV-2 pandemic demonstrated the human and economic toll that can accompany the spillover and spread of a zoonotic disease in humans. Although the success of vaccine development efforts in response to the pandemic were a triumph of modern vaccinology, SARS-CoV-2 remains the only coronavirus for which licensed vaccines are available. To date, nine coronaviruses are known to infect humans, three of which have caused epidemics or pandemics in the last 20 years and two of which have been identified in humans in only the past two years, underscoring that more coronaviruses than previously appreciated currently circulate in humans and pose zoonotic threats (*1–4*). Developing vaccines for additional coronaviruses, both known and unknown, is therefore a public health priority (*5*).

Among the known human-infecting coronaviruses, MERS-CoV stands out due to its high case fatality rate, estimated at 35% (*6*, *7*). Since its discovery in 2012, MERS-CoV infections have been reported in 27 countries, mostly from contact with dromedary camels, although sporadic human-to-human transmission has also occurred (*8*). Beyond its immediate value in the prevention of severe respiratory disease and death caused by MERS-CoV infection, a safe and effective MERS-CoV vaccine would provide a foundation for developing broadly protective vaccines that could prevent zoonotic spillover of known or unknown members of the merbecovirus subgenus. Three MERS-CoV vaccine candidates have entered clinical trials, although none have advanced to licensure (*9–11*).

As the major surface antigen and the target of neutralizing antibodies, the spike (S) protein has been the focus of most MERS-CoV vaccine development efforts. S is a large trimeric class I viral fusion protein that is cleaved into two subunits, S_1_ and S_2_ (*12*). S_1_ contains the N-terminal domain (NTD) and receptor binding domain (RBD) that mediate virus attachment to host cells by binding sialosides and the proteinaceous receptor dipeptidyl peptidase 4 (DPP4), respectively (*13–17*). S_2_ comprises the fusion machinery that merges the virus and host membranes. Several studies have shown that the majority of serum neutralizing activity after infection or immunization, as well as the most potently neutralizing monoclonal antibodies, target the RBD and NTD (*18–25*), although neutralizing and protective antibodies targeting S_2_ have also been characterized (*26–34*). Immunization with the MERS-CoV RBD, S_1_, S_2_, full-length S, and prefusion-stabilized S ectodomain have been evaluated in preclinical animal models and found to induce robust antibody responses and in some cases protection against challenge (*18*, *19*, *22*, *23*, *26*, *28*, *35–40*). Nevertheless, new vaccine design technologies and lessons from SARS-CoV-2 vaccine development efforts may allow the generation of vaccine candidates with improved safety, immunogenicity, and manufacturability.

A major lesson of the recent SARS-CoV-2 pandemic was that pre-existing platform technologies were essential to rapid vaccine development. For example, the prefusion-stabilizing 2P mutations that were identified using MERS-CoV as a prototype pathogen in 2017 (*38*) enabled essentially immediate structure determination and vaccine design based on the SARS-CoV-2 Spike (*40–42*). The readiness of mRNA as a rapid response vaccine platform also proved critical (*43*, *44*). Additional platform technologies were clinically de-risked during the pandemic, including computationally designed two-component protein nanoparticle vaccines (*45*, *46*). Over the last several years, our groups and others have shown that these immunogens elicit potent neutralizing antibody responses against a number of viral pathogens by efficiently trafficking to lymph nodes and enhancing B cell activation (*37*, *47–58*). In response to the SARS-CoV-2 pandemic, we developed a nanoparticle vaccine that displays 60 copies of the SARS-CoV-2 RBD on the icosahedral nanoparticle I53-50 (*59*). RBD-I53-50 induced robust neutralizing antibody responses in mice and NHPs and was found to be safe and immunogenic in clinical trials, leading to its licensure in multiple jurisdictions under the name SKYCovione™ (*45*, *60–63*). Collectively, these results suggest that two-component nanoparticles may be a robust and versatile vaccine platform.

Here, we generate two-component protein nanoparticle immunogens of diverse geometries displaying the MERS-CoV S-2P trimer, RBD, or NTD that elicit potent neutralizing antibody responses targeting antigenic sites in the S_1_ subunit. Our findings identify several promising vaccine candidates for further development and establish two-component nanoparticles as a generalizable platform for betacoronavirus vaccine development.

## Results

### Production and characterization of nanoparticle immunogens

Based on our successful previous application of two-component self-assembling nanoparticles to the development of sarbecovirus vaccines (*37*, *45*, *62–64*), we set out to evaluate a series of nanoparticle immunogens for MERS-CoV. The prefusion-stabilized MERS S-2P trimer was previously shown to elicit neutralizing antibodies in mice as a soluble trimer (*38*), when displayed on nanoparticles (*37*), or when delivered by mRNA (*40*). To further explore the immunogenicity of this antigen, we displayed 4 or 20 copies of the S-2P trimer on two distinct nanoparticles, T33_dn10 and I53-50. T33_dn10 is composed of four copies each of two different designed trimeric proteins that assemble into a 24-subunit nanoparticle with tetrahedral symmetry (*65*), while I53-50 is constructed from 12 pentamers and 20 trimers that form a 120-subunit complex with icosahedral symmetry (*59*). In each case, the S-2P ectodomain trimer with a C-terminal foldon was genetically fused to a trimeric component with externally oriented N termini (**Fig. 1A** and **Table S1**). Given the success of our RBD-I53-50 nanoparticle vaccine against SARS-CoV-2 (*45*, *64*) and the key role of the NTD in the attachment of MERS-CoV to cells (*16*, *17*), we also evaluated I53-50-based nanoparticle immunogens displaying the MERS-CoV RBD or NTD. We genetically fused these domain-based antigens to the N terminus of the I53-50A trimer to enable display of 60 copies on the I53-50 nanoparticle in a manner analogous to our previously described SARS-CoV-2 RBD nanoparticle vaccine (*64*).

**Fig. 1.**
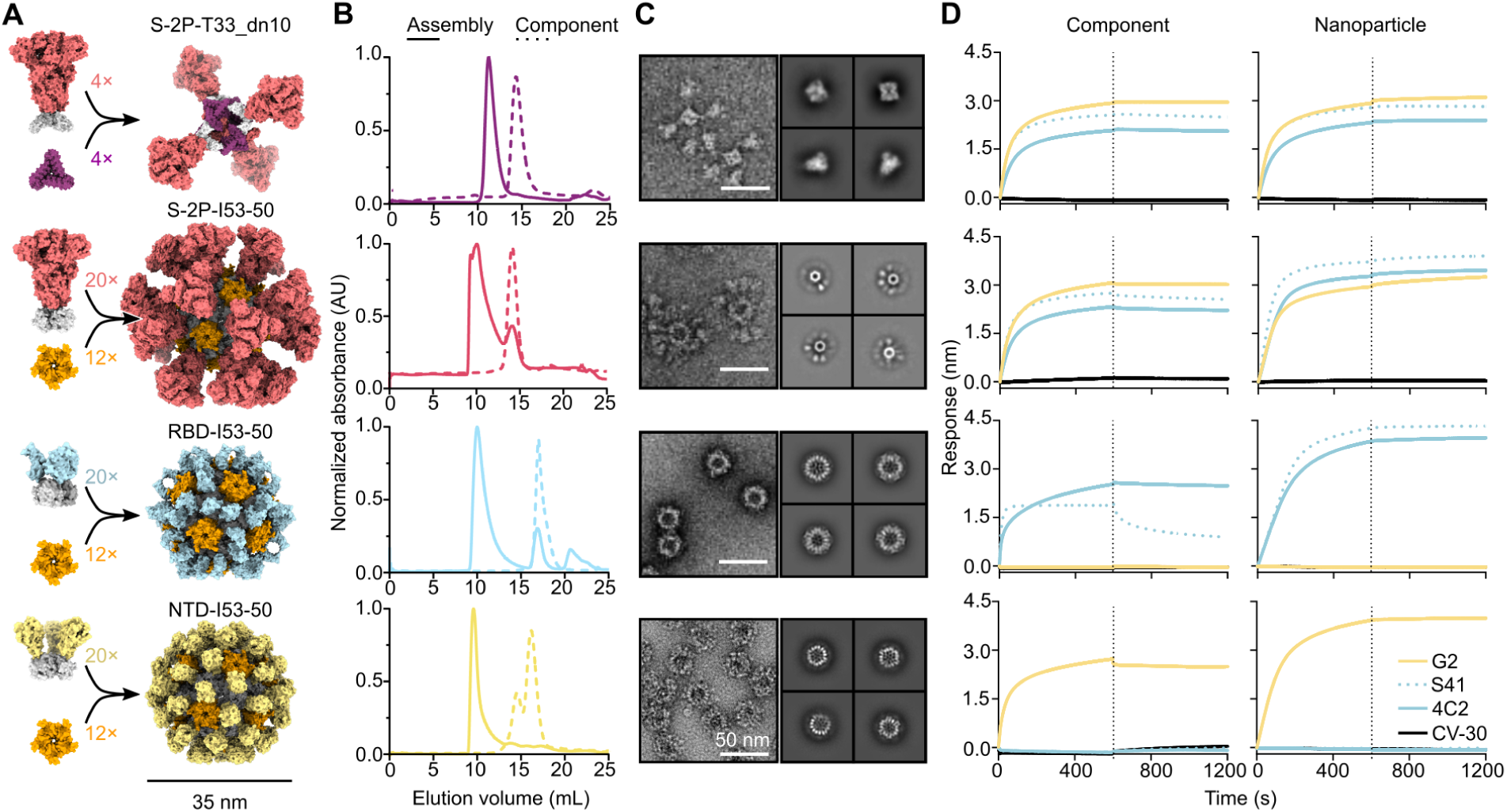
Production and characterization of nanoparticle immunogens. (**A**) Schematics of MERS-CoV nanoparticle immunogen assembly. (**B**) Size exclusion chromatograms of MERS-CoV nanoparticle components (dotted lines) and assembled nanoparticles (solid lines) on a Superose 6 10/300 GL. (**C**) *Left,* raw micrographs and *right,* 2D class averages of negatively stained SEC-purified nanoparticles. For S-2P-T33_dn10, class averages of the nanoparticle core and displayed S-2P trimer are shown on top and bottom, respectively. Scale bar, 50 nm. (**D**) Binding of *left*, antigen-bearing components prior to assembly and *right*, SEC-purified nanoparticle immunogens to mAbs G2 (NTD-specific), S41 (RBD-specific), 4C2 (RBD-specific), and CV-30 (SARS-CoV-2 S RBD-specific; negative control).

We secreted these glycoprotein antigen-bearing components from transfected Expi293F cells and purified them from clarified supernatants using immobilized metal affinity chromatography. We also expressed and purified the soluble MERS-CoV S-2P trimer as a fusion to the foldon trimerization domain for use as a benchmark. Size exclusion chromatography (SEC) of the S-2P-T33_dn10A and S-2P-I53-50A trimers showed that these components eluted at the same volume as the S-2P foldon trimer, while the RBD– and NTD-I53-50A components eluted later, as expected due to their smaller size (**Fig. 1B** and **Fig. S1A**). After mixing each antigen-bearing component with the appropriate second component (i.e., trimeric T33_dn10A or pentameric I53-50B), a shift to an earlier SEC elution volume was observed, suggesting nanoparticle assembly. Small amounts of residual unassembled components were observed in the S-2P– and RBD-I53-50 assembly reactions, as expected given that each contained a slight excess of the trimeric component (see Methods). Analysis of the SEC-purified nanoparticles by SDS-PAGE confirmed that each preparation contained both of the expected protein components, and dynamic light scattering (DLS) indicated monodisperse nanoparticles with the expected hydrodynamic diameters (**Fig. S1, B and C**). Negative stain electron microscopy (nsEM) supported the SEC and DLS data, revealing fields of monodisperse nanoparticles of the expected sizes and shapes. S-2P-T33_dn10 clearly displayed four copies of the S-2P trimer, although these were flexibly linked to the underlying nanoparticle core: we obtained separate classes during 2D averaging corresponding to the tetrahedral nanoparticle and the displayed prefusion S trimer (**Fig. 1C**). nsEM of S-2P-I53-50 revealed spherical assemblies densely decorated with Spikes that resembled micrographs previously obtained for sarbecovirus S-I53-50 immunogens (*66–68*). RBD– and NTD-I53-50 also formed well-defined nanoparticles with additional densities on the nanoparticle surface, although these were less clearly defined than displayed S-2P due to the smaller size of the RBD and NTD antigens. Having confirmed expression of the antigen-bearing components and assembly of the nanoparticle immunogens, we next sought to evaluate retention of antigenicity using bio-layer interferometry (BLI) and a panel of conformation-specific monoclonal antibodies (mAbs) (**Fig. 1D**). As expected, the two RBD-specific antibodies S41 (*69*) and 4C2 (*22*) bound to all antigen-bearing components and nanoparticles except for NTD-I53-50A and NTD-I53-50. Conversely, the NTD-directed antibody G2 (*18*, *25*) bound to all constructs except RBD-I53-50A and RBD-I53-50. Dissociation was unambiguously observed only for the RBD-I53-50A–S41 complex; as expected, the highly avid interactions between the corresponding RBD-I53-50 nanoparticle and the S41 mAb prevented dissociation. Our negative control antibody CV-30, a SARS-CoV-2 S RBD-directed neutralizing antibody isolated from a convalescent patient (*70*), did not bind to any of our MERS constructs. Together, these data demonstrate the production of a series of well-defined and antigenically intact nanoparticle immunogens for MERS-CoV displaying the prefusion-stabilized S-2P trimer, RBD, or NTD.

### Antibody responses elicited by MERS-CoV nanoparticle immunogens in mice

We evaluated the immunogenicity of our MERS-CoV nanoparticles by measuring antigen-specific and neutralizing antibody titers in serum after immunizing groups of 10 BALB/c mice twice with AddaVax-adjuvanted immunogens four weeks apart (**Fig. 2A**). Two weeks post-prime, the highest levels of S-2P-specific antibodies were observed in the sera of animals that received RBD-I53-50 (geometric mean titer (GMT) EC_50_ of 5.6×10^3^), followed by S-2P-I53-50 (1.6×10^3^); all other groups had GMTs <10^3^ (**Fig. 2B** and **Fig. S2A**). Two weeks post-boost, all groups but the bare I53-50 nanoparticle control showed high antibody titers, ranging from 3.2–8.4×10^4^, that slightly waned by week 8 (7.0×10^3^–1.9×10^4^). S-2P and S-2P-T33_dn10 elicited cross-reactive serum antibodies that bound to SARS-CoV and SARS-CoV-2 HexaPro trimers (*71*), but we did not observe similar cross-reactivity from the nanoparticles displaying the domain-based antigens (**Fig. 2C** and **Fig. S2B**). S-2P-I53-50 sporadically elicited low levels of cross-reactive antibodies, perhaps indicating that conserved epitopes in the S_2_ subunit are inaccessible to B cells when densely displayed on I53-50. No detectable binding was observed against the S-2P trimer from the embecovirus OC43 in any sera. Serum neutralizing activity against a panel of VSV pseudotypes bearing closely related MERS-CoV Spikes (EMC, London, Kenya, and South Korea; the latter three have 99% amino acid identity to the vaccine strain; **Table S1**) largely mirrored the binding antibody titers. Specifically, RBD-I53-50 was the only immunogen that elicited appreciable neutralizing activity post-prime (EMC strain GMT 1.1×10^2^ and South Korea strain GMT 1.5×10^2^), while potent neutralization was observed post-boost for all groups at week 6, with subsequent waning by week 8 (**Fig. 2D** and **Fig. S3**). Slightly more separation between the groups was observed in the post-boost neutralization data than in the antigen-specific antibody titers. Specifically, although all groups elicited robust neutralizing activity, the soluble S-2P trimer and S-2P-T33_dn10 generally induced the highest titers, while the nanoparticles displaying domain-based antigens were in some cases lower.

**Fig. 2.**
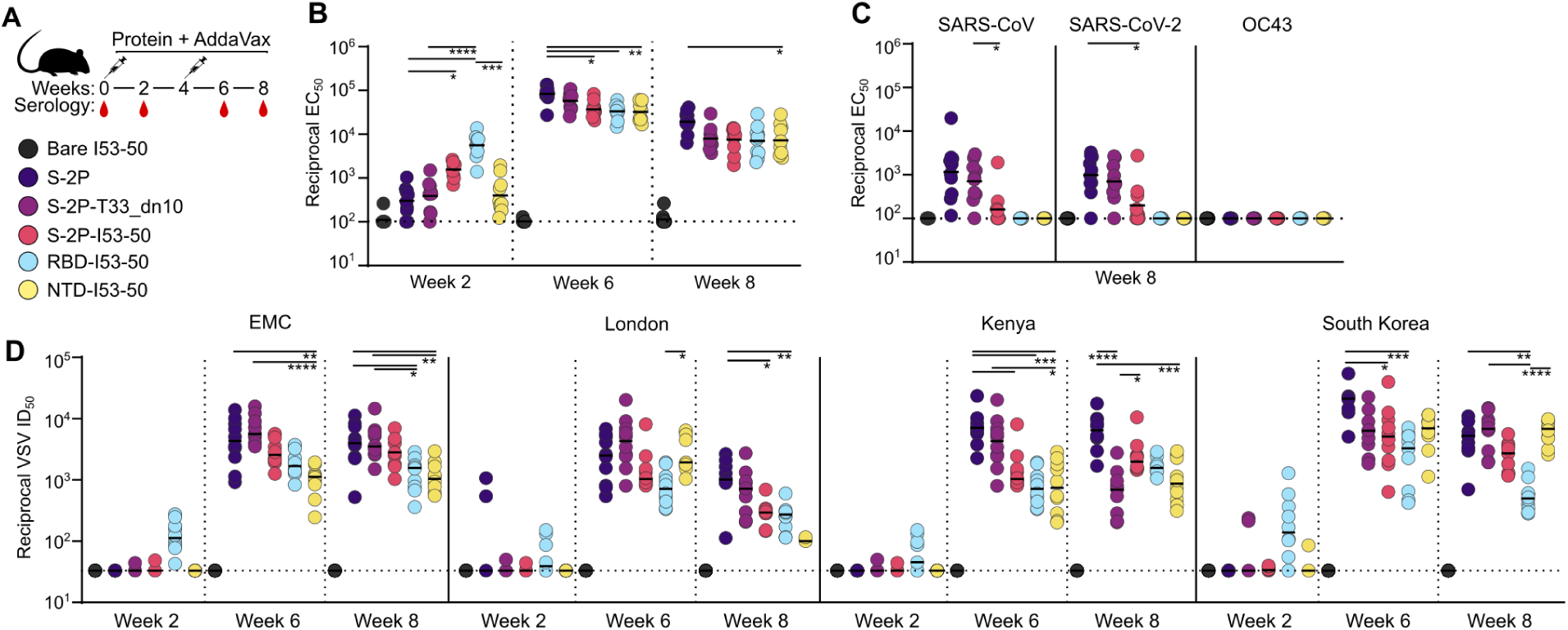
Antibody responses elicited by MERS-CoV nanoparticle immunogens in mice. (**A**) Study design and groups. Groups of 10 BALB/c mice were immunized at weeks 0 and 4, and serum was obtained at weeks 0, 2, 6, and 8. (**B**) Serum antibody binding titers against vaccine-matched (EMC) MERS-CoV S-2P, measured by ELISA. (**C**) Serum antibody titers against SARS-CoV HexaPro, SARS-CoV-2 HexaPro, and OC43 S-2P. (**D**) Vaccine-elicited neutralizing activity against VSV pseudotyped with closely related MERS-CoV EMC, London variant, Kenya variant, or South Korea variant spikes. Groups were compared using Kruskal-Wallis followed by Dunn’s multiple comparisons. ****, p < 0.0001; ***, p < 0.001; **, p < 0.01; *, p < 0.1. The mouse immunization study was performed twice, and representative data from one study are shown.

### Epitope mapping of vaccine-elicited antibodies

We used serum competition assays to map the specificity of the antibody responses elicited by the S-2P trimer and nanoparticle immunogens. We measured the week 8 serum dilutions required to compete with hDPP4 and G2 binding to examine receptor-blocking and NTD-directed responses, respectively (**Fig. 3A and S4**). We observed comparable hDPP4 competition across the S-2P, S-2P-T33_dn10, S-2P-I53-50, and RBD-I53-50 groups, suggesting that all of these vaccines induced similar levels of receptor-blocking antibodies. The bare I53-50 nanoparticle control group did not elicit detectable hDPP4– or G2-competing antibodies as expected. We observed more variation among immunogens in the competition of our polyclonal sera with G2. Although none of the sera completely blocked G2 binding under the conditions tested, they could be stratified into three groups. Sera from mice that received NTD-I53-50 and S-2P-T33_dn10 most potently competed with G2 binding, followed by the S-2P trimer and S-2P-I53-50, followed by the RBD-I53-50 nanoparticle. The competition from RBD-I53-50-induced antibodies, albeit weak, was consistent across the sera from all mice (**Fig. S3D**). We interpret these data to indicate that some RBD-directed antibodies sterically compete with the binding of NTD-directed antibodies, a phenomenon that is similar to blockade of DPP4 binding by some NTD-directed antibodies (*23*, *25*).

**Fig. 3.**
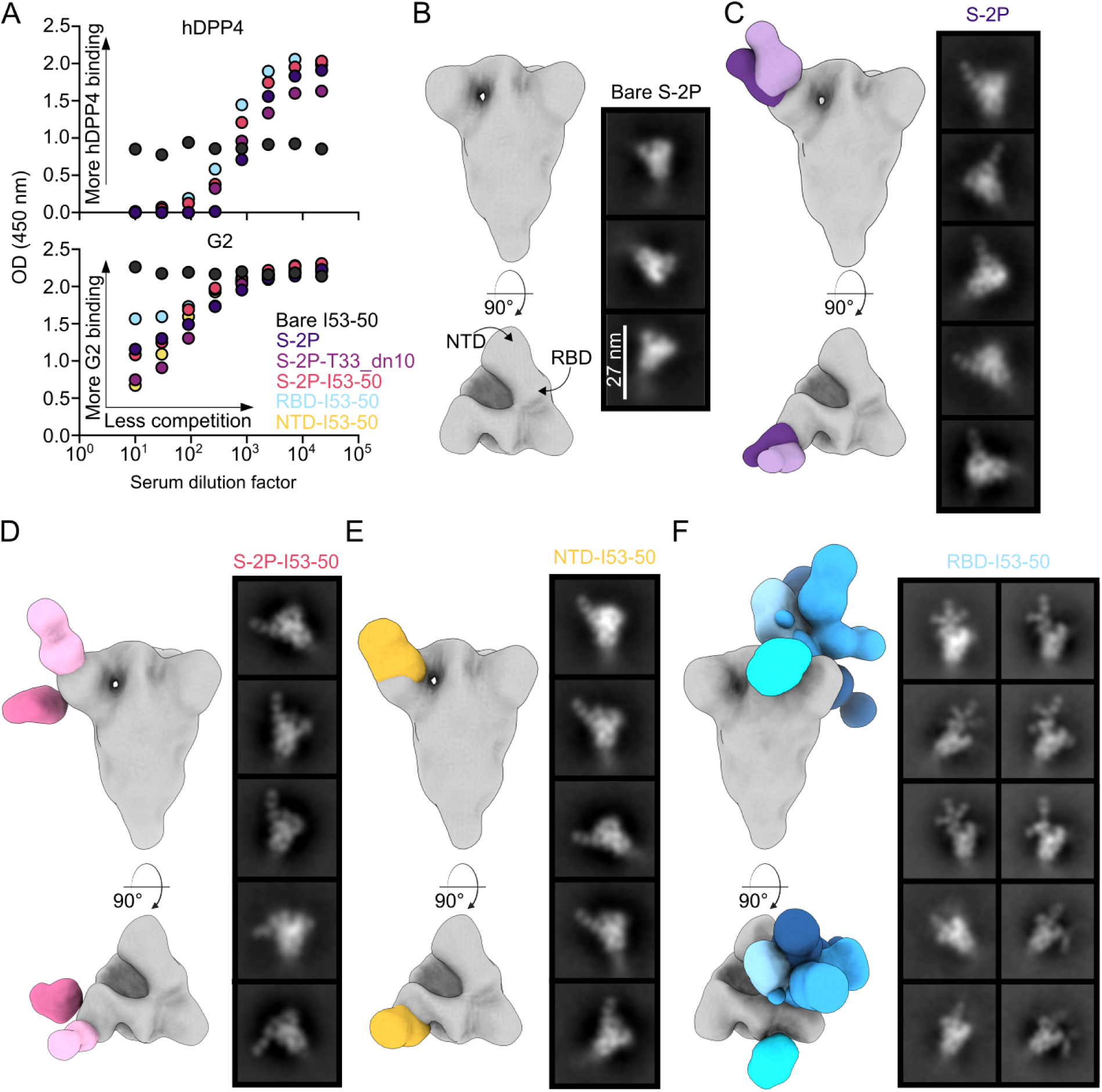
Epitope mapping of vaccine-elicited antibodies. (**A**) Serum competition of study groups with hDPP4 (top) and G2 (bottom) against MERS-CoV S-2P. (**B**) 3D reconstructions of MERS S-2P with no bound Fabs and selected 2D class averages. Scale bar, 27 nm. (**C** to **F**) Representative reconstructions from week 8 sera from mice immunized with S-2P (**C**), S-2P-I53-50 (**D**), NTD-I53-50 (**E**), and RBD-I53-50 (**F**).

We next visualized the dominant serum antibody specificities elicited by the S-2P trimer and the three I53-50 nanoparticle immunogens using nsEM polyclonal epitope mapping (ns-EMPEM) (*72*, *73*). To do this, we first generated a low-resolution reconstruction of the MERS-CoV S-2P trimer, which was consistent with structures obtained in previous reports (**Fig. 3B**) (*17*, *38*, *73*). We then pooled and processed mouse sera from week 10 to generate polyclonal Fab complexes with MERS-CoV S-2P and mapped the densities obtained from ns-EMPEM onto the S-2P trimer reconstruction. In sera from mice immunized with soluble S-2P, we observed two slightly distinct classes of NTD-directed Fabs that bind the antigenic site targeted by G2 and many NTD-directed neutralizing antibodies against SARS-CoV-2 (**Fig. 3C**) (*25*, *74–76*). We did not detect RBD-directed Fab classes using these sera, possibly due to spike triggering and unfolding (*77*, *78*). We also only observed NTD-directed Fab classes in the sera of mice that received S-2P-I53-50 (**Fig. 3D**). One of the two dominant classes targets the same antigenic site at the apex of the NTD, while the second class binds the side of the NTD with an almost perpendicular angle of approach. Interestingly, these two classes recapitulate the specificities of cross-reactive antibodies against MERS-CoV S-2P identified in a recent study after immunization with a two-component nanoparticle immunogen displaying the SARS-CoV S-2P trimer (*37*).

As expected, the domain-based NTD– and RBD-I53-50 nanoparticle immunogens elicited antibodies that bind each respective domain. All of the Fab-antigen complexes observed in the anti-NTD-I53-50 sera target the antigenic site at the apex of the NTD; we did not see 2D classes showing Fabs bound to the side of the NTD as we did for S-2P-I53-50 (**Fig. 3E**). By contrast, in sera from mice immunized with RBD-I53-50 we observed a number of distinct classes of antibodies targeting various epitopes (**Fig. 3F**). Strikingly, several 2D class averages clearly showed multiple Fabs simultaneously bound to the same RBD, indicating that they bind non-overlapping epitopes.

Our inability to resolve RBD-directed densities from the soluble S-2P and S-2P-I53-50 groups was surprising given the apparent immunodominance of the RBD in MERS-CoV and other coronaviruses (Addetia et al. and (*79–81*)). Furthermore, our hDPP4 competition data clearly indicate the presence of high levels of RBD-directed antibodies in these sera (**Fig. 3A**). We note that we had to optimize our experimental conditions to prevent destabilization of the Spike protein (*82*) by shortening the polyclonal Fab-antigen incubation time and using a lower molar ratio of polyclonal Fabs to antigen (see Methods). It is likely that these conditions prevented us from observing all Fab classes prevalent in the sera. As a result, we only provide qualitative interpretations of our ns-EMPEM data. Nevertheless, taken together our epitope mapping data suggest that the RBD and NTD of the MERS-CoV spike are immunodominant during vaccination, as was observed in many studies of antibody responses to SARS-CoV-2 (*80*, *83–86*) and a limited number of studies of other betacoronaviruses (*73*, *81*, *84*, *85*). Our results further suggest that a single antigenic site is immunodominant in the NTD, whereas several antigenic sites in the RBD are highly immunogenic.

### Protection against challenge with mouse-adapted MERS-CoV

Finally, we evaluated the protection afforded by each immunogen in a mouse immunogenicity and challenge study. We immunized 288/330^+/+^ mice that express a chimeric DPP4 receptor in the C57BL/6J genetic background with two doses of our S-2P or RBD-based immunogens and subsequently challenged them at study week 15 with 1×10^5^ pfu of mouse-adapted MERS-CoV (maM35c4) (*87*, *88*) (**Fig. 4A**). This model has previously been shown to recapitulate many aspects of the acute respiratory distress syndrome that humans experience upon MERS-CoV infection (*87*, *88*). The levels of S-2P-specific antibodies in sera collected two weeks post-prime and post-boost were generally similar to those observed in our earlier study in BALB/c mice, except that the post-prime titers were slightly higher for the S-2P-based immunogens in this study (**Fig. 4B**). Similar results were obtained in pseudovirus neutralization assays: the data mirrored our earlier BALB/c immunogenicity study except that the S-2P-displaying nanoparticles induced detectable neutralizing activity at week 2 in addition to the RBD-I53-50 nanoparticle (**Fig. 4C**). Overall, the trends in antigen-specific and neutralizing antibody titers between the 288/330^+/+^ and BALB/c mice were comparable, showing slightly higher responses from the nanoparticle groups after priming and similar responses after boosting.

**Fig. 4.**
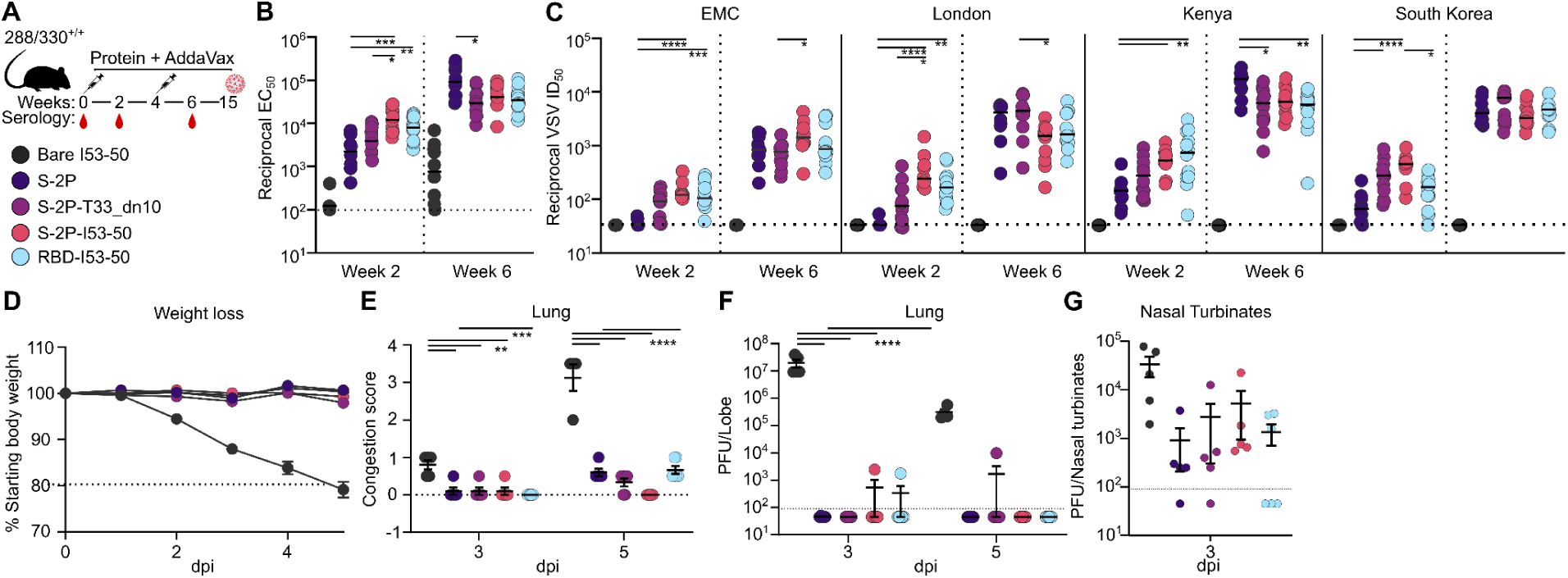
Protection against challenge with mouse-adapted MERS-CoV. (**A**) Challenge study design and groups. Groups of 288/330^+/+^ mice were immunized at weeks 0 and 4, serum was obtained at weeks 0, 2, and 6, and the mice were challenged at week 15. (**B**) Serum antibody binding titers against vaccine-matched (EMC) MERS-CoV S-2P, measured by ELISA. (**C**) Vaccine-elicited neutralizing activity against VSV pseudotyped with closely related MERS-CoV EMC, London variant, Kenya variant, or South Korea variant spikes. Groups were compared using Kruskal-Wallis followed by Dunn’s multiple comparisons. ****, p < 0.0001; ***, p < 0.001; **, p < 0.01; *, p < 0.1. (**D**) Changes in body weight after maM35c4 challenge. (**E**) Lung congestion score. Data analyzed with two-way ANOVA or mixed model followed by Sidak’s multiple comparison. ****, p < 0.0001; *** p < 0.001; ** p < 0.01. (**F**) Viral titers in the lungs of challenged mice from (3 dpi: n = 5 for bare I53-50, S-2P, S-2P-T33_dn10, and S-2P-I53-50; n = 6 for RBD-I53-50. 5 dpi: n = 4 for bare I53-50; n = 5 for S-2P and S-2P-I53-50; n = 6 for S-2P-T33_dn10 and RBD-I53-50. Data were analyzed with two-way ANOVA or mixed model followed by Dunnett’s multiple comparison. ****, p < 0.0001. (**G**) Viral titers in nasal turbinates of challenged mice. n = 5 for bare I53-50, S-2P, S-2P-T33_dn10, and S-2P-I53-50; n = 6 for RBD-I53-50. Data analyzed with one-way ANOVA with Geisser-Greenhouse correction followed by Dunnett’s multiple comparison.

maM35c4 challenge at week 15 allowed us to evaluate the efficacy of our immunogens against the development of severe disease. All 11 control mice immunized with two doses of bare I53-50 lost 20% of their starting weight by day 5 post-infection and had to be euthanized (**Fig. 4D**). By contrast, all mice given two doses of MERS-CoV immunogens were protected from severe disease, showing no appreciable weight loss after infection. Furthermore, the mice receiving MERS immunogens displayed undetectable to mild lung damage 3 and 5 days post-infection, while those receiving the bare I53-50 control immunogen showed considerable damage at day 5 (**Fig. 4E**). While viral titers in the lung were below the limit of detection in nearly all mice immunized with antigen-bearing immunogens throughout the duration of the experiment, viral titers in nasal turbinates were reduced ≥1-2 logs relative to those in the bare I53-50 control group at 3 days post-infection (**Fig. 4F,G**). Together, these data show that immunization with prefusion-stabilized S-2P and a series of two-component nanoparticle immunogens elicited potent neutralizing antibody responses and protected 288/330^+/+^ mice from developing severe disease when challenged with a lethal dose of maM35c4.

## Discussion

Here we demonstrated that two-component nanoparticle vaccines elicit potent neutralizing activity and protective responses against a merbecovirus. The successful application of two-component nanoparticle vaccines to a second subgenus of betacoronaviruses suggests that their utility may extend even further across the *Coronaviridae*. Although several other protein nanoparticle technologies were evaluated as SARS-CoV-2 vaccines (*89–96*) and some have begun to be applied to other betacoronaviruses (*97–99*), to our knowledge NVX-CoV2373 from NovaVax and the RBD-I53-50 nanoparticle vaccine SKYCovione™ are the only two that have been licensed. Two-component protein nanoparticle vaccine candidates for RSV and RSV/HMPV have also been found to be safe and immunogenic in clinical trials (*48*, *100–102*), and a two-component mosaic nanoparticle vaccine for influenza virus is currently in Phase I clinical trials ((*54*); NCT04896086 and NCT05968989). These precedents combine with the data presented here to establish two-component protein nanoparticles as a clinically de-risked platform for pandemic preparedness vaccine development.

The SARS-CoV-2 pandemic also led to a uniquely deep understanding of the mechanisms of antibody-mediated protection against a betacoronavirus. For SARS-CoV-2, although effector functions from non-neutralizing antibodies can also protect (*30*, *103–107*), the most potently neutralizing antibodies target the RBD and a single immunodominant antigenic site in the NTD (*74*, *76*, *80*, *108*, *109*). Our data extend those observations to merbecoviruses. We found that all of the vaccine candidates tested, including those based on the prefusion-stabilized S-2P trimer or nanoparticles displaying domain-based antigens, elicit robust neutralizing activity. The MERS-CoV RBD-I53-50 nanoparticle elicited the most potent antibody responses after a single immunization, mirroring results we previously obtained for SARS-CoV-2 (*64*). However, unlike that study, the MERS-CoV S-2P-based immunogens induced higher neutralizing activity post-boost. It is possible that this difference may derive in part from higher physical stability of the prefusion-stabilized MERS-CoV Spike compared to SARS-CoV-2 S-2P. This notion is consistent with the enhanced immunogenicity of the more stable HexaPro variant of the SARS-CoV-2 Spike (*71*, *110*) and stabilized variants of prefusion RSV F and HIV Env (*111*, *112*). It is also possible that neutralizing antibodies targeting both the RBD and NTD, rather than just one of these domains, account for the potent neutralizing activity elicited by the S-2P-based immunogens (*18*). In an accompanying manuscript we show that most of the MERS-CoV neutralizing activity in the sera of infected humans targets the RBD and NTD (Addetia et al.), and our results obtained with NTD-I53-50 clearly establish the neutralizing activity of vaccine-elicited antibodies against the NTD. These antibodies, like several previously described mAbs (*18*, *19*, *23*, *113*, *114*), appear to primarily target a dominant antigenic site near the α-2,3-linked sialoside binding site (*17*, *25*). The role of sialoside binding in MERS-CoV attachment to host cells (*16*) suggests a potential mechanism for neutralization by NTD-directed antibodies beyond blocking DPP4 engagement sterically. However, given the rapid evolution of the SARS-CoV-2 NTD during the pandemic (*76*, *115–117*) and the high sequence diversity in merbecovirus NTDs, it is not clear that an NTD-based immunogen would be preferred for further development. By contrast, antibodies elicited by MERS-CoV RBD-I53-50 target a number of distinct, non-overlapping epitopes in the RBD, suggesting that this vaccine candidate may neutralize or protect against MERS-CoV variants with some antigenic variation in the RBD, as was observed for RBD-based nanoparticle immunogens for SARS-CoV-2 (*63*, *93*, *94*, *97*, *118*, *119*).

In summary, we found that the prefusion-stabilized MERS-CoV Spike and several two-component nanoparticle immunogens induce robust and protective immune responses against MERS-CoV in vaccinated mice. The high performance of the RBD-I53-50 nanoparticle and its use of the same platform as the licensed SARS-CoV-2 vaccine SKYCovione™ motivates further development of this vaccine candidate. More generally, our data highlight two-component protein nanoparticle immunogens as a platform technology that may be broadly useful for designing vaccines against betacoronaviruses and other virus families.

## Acknowledgements

The authors thank Sebastian Ols, Kenneth Carr, and Andrew Borst for helpful discussions.

## Author contributions

C.W.C., M.C.M., A.C.W., D.V., and N.P.K. designed the study. C.W.C., M.C.M, and A.C.W. designed and characterized immunogens. C.W.C. and K.R.S. performed *in vitro* experiments, and N.J.C. performed challenge experiments. C.W.C., M.C.M., A.A., C.S., J.T.B., A.D., A.V., R.R., and G.G.H. produced proteins and provided reagents. C.T., I.W., E.M.L., and M.H. conducted and performed animal immunizations. M.A., C.D., and A.H. provided and maintained cells. C.W.C. and K.R.S. analyzed the data. A.T.M., A.S., and R.S.B. provided oversight of immunization and challenge studies. C.W.C., N.P.K., D.V., K.R.S., E.M.L., and A.S. wrote the manuscript with input from all authors.

## Materials and Methods

### Protein purification and characterization

All constructs contained a C-terminal histidine affinity tag and were codon optimized by GenScript for mammalian cell expression. Expi293F cells were transiently transfected using PEI MAX and cultured for three days 37°C. Cells were harvested, centrifuged, and filtered to obtain clarified supernatants. Clarified cell supernatants were buffered with 5 mL of 5M NaCl and 7 mL of Tris pH 8.0 per 100 mL of supernatant and batch bound with Ni Sepharose Excel resin while shaking for 30 minutes to an hour at room temperature. Resin were collected in a gravity column, washed with five column volumes of wash buffer containing 25 mM Tris pH 8.0, 150 mM NaCl, and 30 mM imidazole, and eluted with three column volumes of elution buffer comprised of 25 mM Tris pH 8.0, 150 mM NaCl, and 300 mM imidazole. Eluted component constructs were further purified using SEC on either a Superose 6 Increase 10/300 gel filtration column if they were spike-bearing, or a Superdex 200 Increase 10/300 gel filtration column if they were domain-bearing constructs. *In vitro* assembly of nanoparticles was conducted using a 1:1 molar ratio of each component and incubated at room temperature for 30 minutes with rocking. Following nanoparticle assembly, all particles were purified one last time through a Superose 6 Increase 10/300 column. All spike and NTD protein components and assembled nanoparticles were purified into 50 mM Tris pH 8.0, 150 mM NaCl, 0.25% w/v Histidine, and 5% glycerol. The RBD component and nanoparticle were purified in 50 mM Tris pH 7.4, 185 mM NaCl, 100 mM Arginine, 4.5% Glycerol, and 0.75% CHAPS.

### Bio-Layer interferometry (BLI)

All samples were diluted in kinetics buffer (0.05% bovine serum albumin and 0.01% Tween 20 in PBS) and measurements were carried out using an Octet Red 96 system at 25°C shaking at 1000 rpm. Antibodies were diluted to a final concentration of 20 ug/mL before being immobilized on ProteinA tips (Sartorius) for 600 s. Unassembled and nanoparticle proteins were diluted to 250 nM and associations were measured for 600 s, followed by dissociation for 900 s in kinetics buffer.

### Immunogenicity Studies

For immunogenicity studies, female BALB/cAnNHsd were purchased from Envigo (order code 047) at 7 weeks of age. Mice were housed in a specific-pathogen free facility within the Department of Comparative Medicine at the University of Washington, Seattle, accredited by the Association for Assessment and Accreditation of Laboratory Animal Care (AAALAC). Animal studies were conducted in accordance with the University of Washington’s Institutional Animal Care and Use Committee. For each immunization, low-endotoxin nanoparticles were diluted to 20 μg/mL in buffer and mixed with 1:1 v/v AddaVax adjuvant (InvivoGen vac-adx-10) to obtain a final dose of 1 μg of immunogen per animal, per injection. At 8 weeks of age, 10 mice per group were injected subcutaneously in the inguinal region with 100 μL of immunogen at weeks 0 and 4. Animals were bled using the submental route at weeks 0, 2, and 6. A terminal bleed was collected at week 8 via cardiac puncture. Whole blood was collected in serum separator tubes (BD #365967) and rested for 30 min at room temperature for coagulation. Tubes were then centrifuged for 10 min at 2,000 *g* and serum was collected and stored at –80°C until use.

### 288/330-hDDP4 transgenic mouse vaccinations and virus challenge experiments

For *in vivo* protection studies, 288/330-hDPP4 transgenic mice were used, which were bred and maintained at the University of North Carolina at Chapel Hill (Animal Welfare Assurance #A3410-01). The study was performed using protocols (#23-085) approved by the UNC Institutional Animal Care and Use Committee (IACUC) and carried out in accordance with the recommendations for care and use of animals by the Office of Laboratory Animal Welfare (OLAW), National Institutes of Health and the Institutional Animal Care. Immunizations were performed in a BSL2 facility accredited by the Association for Assessment and Accreditation of Laboratory Animal Care (AAALAC). The subsequent challenge study was conducted in a BSL3 facility at the University of North Carolina. Briefly, groups of 288/330-hDPP4 transgenic mice (n=5/6 per condition and time point, mixed sexes, ∼19–23 weeks of age) were vaccinated subcutaneously in the inguinal region with 100 μL of immunogen with 1 μg of nanoparticle immunogen at weeks 0 and 4. Animals were bled via submandibular route at –1, 2, and 6 weeks. Mice were then moved into the BSL3 and acclimated for a few days. For infection, mice were anesthetized with an intraperitoneal delivery of xylazine and ketamine and intranasally inoculated with 1×10^5^ PFU of MERS-CoV maM35c4. Mice were monitored daily for weight loss and mortality. On 3 dpi and 5 dpi, groups of mice were euthanized with isoflurane and nasal turbinates and the caudal lobe of the right lung were collected to determine viral load by plaque assay. For plaque assay, harvested tissues were homogenized in 1× PBS, and the resulting homogenate was serial-diluted onto a confluent monolayer of Vero E6 cells, followed by agarose overlay. Plaques were then visualized with an overlay of Neutral Red on 3 dpi.

### Enzyme-linked immunosorbent assay (ELISA)

Costar 96-well high protein binding plates were coated overnight at 4°C with spike proteins at 2 ug/mL and blocked in 200 uL of blocking buffer composed of (TBST: 1x Tris-buffered saline with 25 mM Tris pH 8.0, 150 mM NaCl, 0.2% Tween 20, and 5% nonfat milk). Plates were then washed 3x with TBST in an automated plate washer (Biotek); all washing steps follow the same protocol. 3-fold serial serum dilutions were made starting at 1:100 and 100 uL were plated and incubated for one hour at room temperature shaking at 500 rpm. Plates were washed then 100 uL of secondary anti-mouse (Cell Signaling Technology) (1:2,000 dilution) or anti-human (SouthernBiotech) (1:5,000 dilution) IgG-HRP were added to each well and incubated shaking at room temperature. Plates were washed before 100 uL per well of TMB were added and developed for 3 minutes, then quenched with 100 uL of 1N HCl. Reading at absorbance at 450 nM was carried out with an Epoch plate reader (Biotek).

### Competition ELISA

Competition ELISAs were performed in a similar manner as the aforementioned ELISA method with minor adjustments. To determine fixed competitor concentrations for 80% maximal binding against S-2P, a 3-fold serial dilution series with a starting dilution of 1:10 was carried out in blocking buffer and 100 uL per well of were applied and incubated on plates coated with S-2P for 45 minutes then washed. 100 uL per well of secondary Goat anti-human IgG-HRP (1:20,000 dilution) were plated for 30 minutes at room temperature. Plates were washed then developed for two minutes with 100 uL of TMB and quenched with 100 uL of 1N HCl. The concentration of competitors at which approximately 80% of maximal binding observed was used for competition assays.

As a positive control for achieving full competition against the competitor, hDPP4-Fc and G2 were biotinylated using EZ-Link™ Sulfo-NHS-LC-Biotin (Thermo Scientific) and 80% maximum binding concentrations against S-2P were determined with biotinylated proteins (hDPP4-Fc-biot and G2-biot). For hDPP4-Fc self-competition, a 3-fold serial dilution was performed starting at 300 ug/ml of hDPP4-Fc in blocking buffer and 100 uL per well were plated on S-2P coated plates for 30 minutes. Plates were washed and 100 uL per well of hDPP4-Fc-biot at 8 ug/mL were added and incubated for 45 minutes. Plates were washed 100 uL per well of secondary Anti-Strep-HRP (1:5,000 dilution) (Thermo Scientific) were plated for 30 minutes. Plates were washed one last time then developed for 2 minutes with TMB and quenched with 1N HCl before reading. For G2 self-competition, a 3-fold serial dilution of G2 starting at 10 ug/mL was carried out and added to S-2P coated plates and G2-biot was added as the competitor at 0.37 ug/mL.

For the sera competition assay, 3-fold serial serum dilutions were performed starting at 1:10 in blocking buffer, transferred to S-2P-coated plates and incubated at room temperature for 30 minutes, then washed. 100 uL per well of hDPP4-Fc or G2 at 1.11 ug/mL or 0.12 ug/mL, respectively, were added and incubated for 45 minutes then washed. Goat anti-human IgG-HRP at a 1:20,000 dilution was added for 30 minutes. Plates were developed for 2 minutes and optical densities were read at 450 nm.

### Generation of pseudoviruses

To produce pseudoviruses for entry and neutralization assays, HEK293T cells were seeded in Dulbecco’s Modified Eagle Medium (DMEM) enriched with 10% Fetal bovine serum (FBS, Hyclone), 1% PenStrep (100 I.U./mL penicillin and 100ug streptomycin) (Gibco 15140-122) at the appropriate density to yield 80% confluency in polylysine-coated 100 mm cell culture dishes and placed in an incubator at 37°C with 5% CO_2_. After 18-22 hr incubation, cells were washed with OPTI-Minimum Essential Media (Opti-MEM, Life Technologies). 24 µg of MERS-CoV-EMC, MERS-CoV-London, MERS-CoV-South Korea, MERS-CoV-Kenya full length S plasmids were prepared in 1.5mL of OPTI-MEM and combined with 60µL of Lipofectamine 2000 (Life Technologies) diluted in 1.5mL of OPTI-MEM and incubated at room temperature for 15-20 min. The mixture was added to the HEK293T cells which were placed for 2 hours in an incubator at 37°C with 5% CO_2_ after which 2mL of DMEM enriched with 20% FBS and 2% PenStrep (200 I.U./mL penicillin and 200ug streptomycin) was added to the transfected cells and incubated overnight. The following day, cells were washed with DMEM and transduced with VSVΔG/Fluc and incubated for 2 hours at 37°C with 5% CO_2_. After washing with DMEM, medium supplemented with anti-VSV-G antibody (I1-mouse hybridoma supernatant diluted to 1:25 from CRL-2700, ATCC) was added to the cells to reduce background from the parental virus and an additional incubation at 37°C with 5% CO_2_ was performed overnight. The next day, the supernatant from the cells was harvested from the 100 mm dishes, further clarified by centrifugation at 3,000xg for 10 minutes, filtered (0.45μm), and concentrated 10 times by using centrifugal devices with 30 kDa cutoff membranes. Pseudoviruses were then aliquoted and frozen at –80°C until used.

### Pseudovirus neutralization assay

VeroE6+TMPRSS 2 cells cultured in DMEM with 10% FBS (Hyclone), 1% PenStrep (100 I.U./mL penicillin and 100 µg streptomycin) and 8ug of puryomycin were subsequently trypsinized, counted and reseeded at ∼40,000 cells per well in cell-culture grade 96 well plates and placed in an incubator at 37°C with 5% CO_2_ overnight. The next day, after cell health and confluency was evaluated, a half-area 96-well plate was prepared with a 1:3 serial dilution of sera in DMEM in 22 μL final volume. 22 μL of pseudovirus was then added to each well and incubated at room temperature for 30-45 min. The media was removed from VeroE6+TMPRSS2 cells and they were subsequently washed 2-3x and 40 μL of the sera/pseudovirus mixture from the half area plate was added to the cells and incubated for 2 h at 37°C with 5% CO_2_ before adding 40 μL of 20% FBS and 2% PenStrep (200 I.U./mL penicillin and 200 µg streptomycin) containing DMEM. Following 18-22 hour incubation, 40 μL of One-GloEX (Promega) was added to the cells and incubated in the dark for 5 min prior to reading on an Agilent BioTek Neo2 plate reader. Relative luciferase units were plotted and normalized in Prism (GraphPad) using a zero value of cells alone and a 100% value of 1:2 virus alone. Nonlinear regression of log(inhibitor) vs. normalized response was used to determine ID_50_ values from curve fits. At least two biological replicates with two distinct batches of pseudovirus were conducted for each serum sample.

### Negative stain electron microscopy (nsEM)

400-mesh carbon coated grids (Electron Microscopy Sciences) were glow-discharged and 3 uL of 20 ug/mL nanoparticles were applied then stained with 2% (w/v) uranyl formate. All nsEM data were collected with a BM-Ceta camera at 57,000x magnification using EPU 2.0 on a 120 kV Talos L120C transmission electron microscope (Thermo Scientific). CTF processing, particle picking, particle extraction, 2D classification, and 3D refinement steps were all performed with CryoSPARC (*127*).

### ns-EMPEM

To obtain Fabs for immune complexes, 500 uL of serum from mice were pooled from the terminal bleed time point to maximize the workable volume for processing. Sera were incubated with Pierce^TM^ Protein G Agarose (Thermo Fisher) at pH 5.0 for 12-24 hours at 4°C and polyclonal antibodies were washed with five column volumes of PBS to remove unbound sera and eluted from gravity filtration columns with three column volumes of 0.1 M glycine, pH 2.5 and neutralized with 1 M Tris pH 8.0 to a final diluted concentration of 50 mM Tris pH 8.0. Eluted IgGs were cleaved with papain in 20 mM NaPO4, pH 6.5, 10 mM EDTA, 20 mM Cysteine at 37°C with shaking for 5-20 hours. Cleaved Fabs were retrieved using gravity filtration columns and dialyzed into PBS before the final step of SEC in a Superdex 200 Increase 10/300 gel filtration column equilibrated with PBS. To prepare immune complexes for ns-EMPEM, purified Fab and MERS S-2P were mixed at a 5:1 ratio and incubated at room temperature with rocking for 30-45 minutes, then immediately applied onto glow-discharged 400-mesh carbon coated grids for ns-EMPEM.

Three rounds of 2D classification were performed, selecting for only spike proteins with clear bound Fab densities. Ab initio 3D reconstruction followed by heterogeneous refinement with no symmetry was used to generate 3D models of immune complexes. Only classes with distinct spike densities that captured a representation of bound Fabs were further processed with 3D refinement. Chimera (*128*) and ChimeraX (*129*) were used to make figures.

## Supplemental Figures

**Fig. S1.**
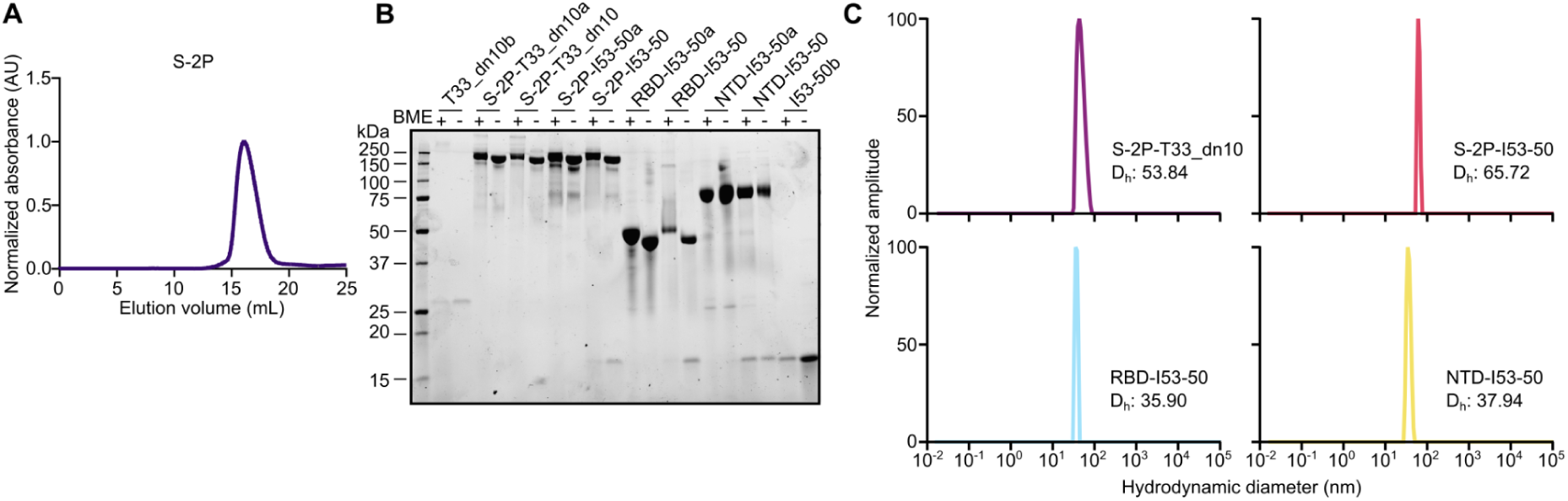
Characterization of MERS immunogens. (**A**) MERS-CoV S-2P SEC on a Superose 6 Increase 10/300 column. (**B**) SDS-PAGE of nanoparticle components before and after assembly under reducing and non-reducing conditions. BME, β-mercaptoethanol. (**C**) DLS of assembled nanoparticles measured on an UNcle (UNchained Labs). Hydrodynamic diameters (D_h_) are indicated.

**Fig. S2.**
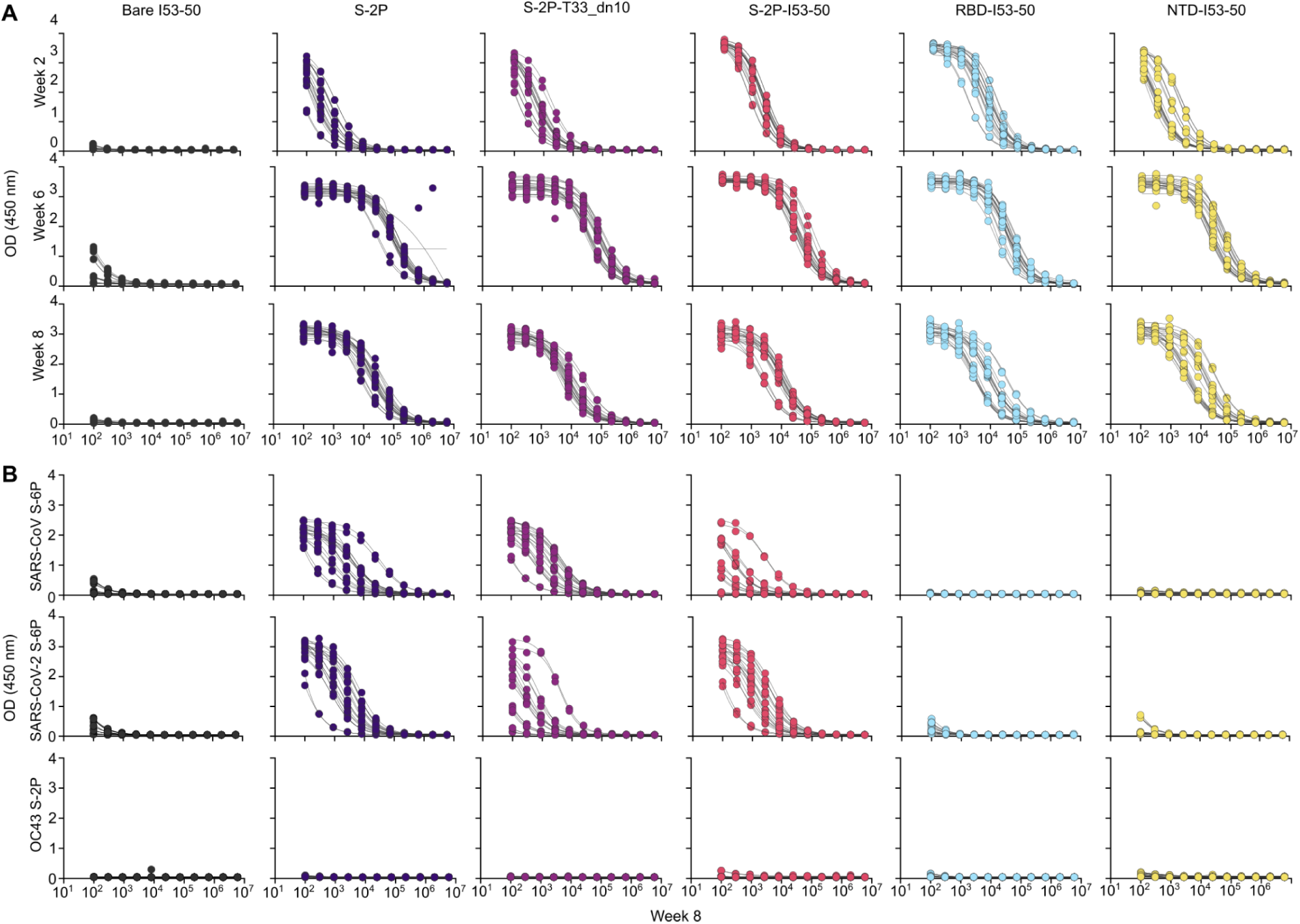
Mouse serum binding to MERS-CoV S-2P, SARS-CoV-2 HexaPro S, and SARS-CoV HexaPro S. (**A**) Raw week 2, 4, and 8 serum ELISA data against vaccine-matched (EMC) MERS-CoV S-2P. (**B**) Raw week 8 serum ELISA data against *top,* SARS-CoV HexaPro; *middle,* SARS-CoV-2 HexaPro; and *bottom,* OC43 S-2P.

**Fig. S3.**
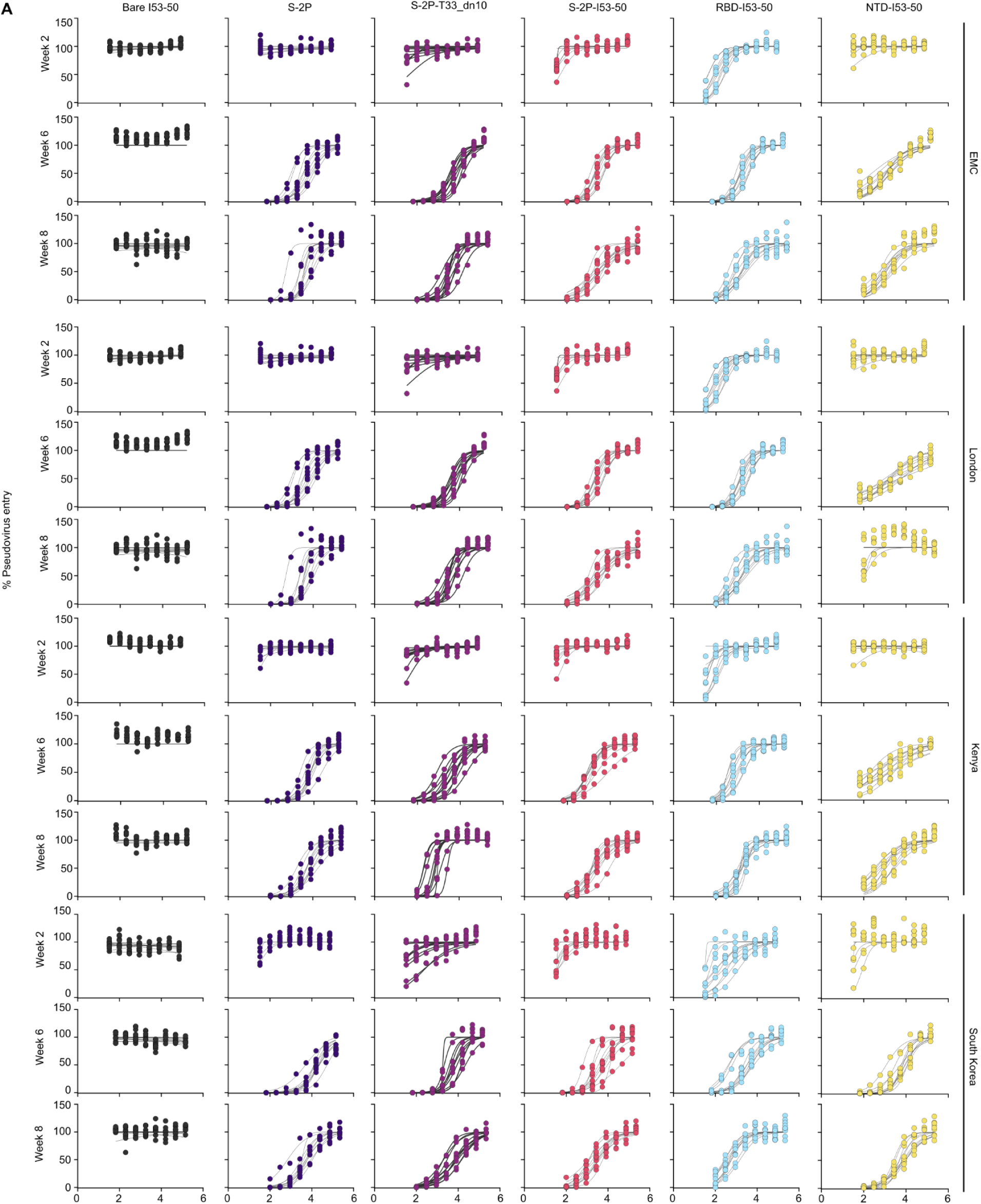
Mouse serum pseudovirus neutralization. **(A)** Raw neutralization data from weeks 2, 6, and 8 using pseudoviruses bearing the MERS-CoV EMC, London, Kenya, and South Korea variant spikes.

**Fig. S4.**
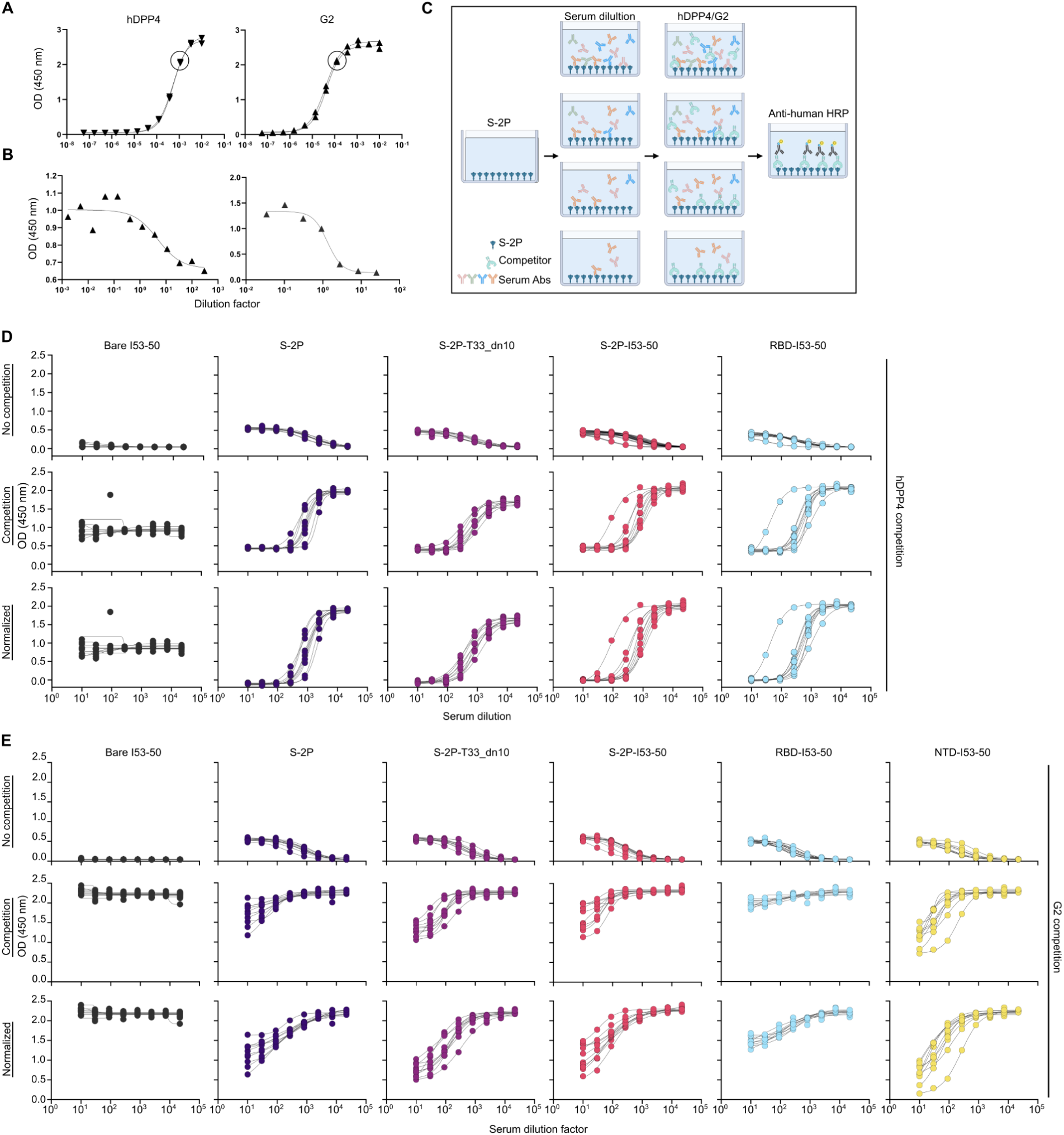
Serum competition against hDPP4 and G2. (**A**) Anti-S-2P ELISA with hDPP4-Fc and G2 to determine concentration of 80% maximum binding (circled). (**B**) Self competition with biotinylated hDPP4-Fc and G2 against S-2P. (**C**) Schematic of serum competition ELISA assay. Serum dilutions were incubated on plates coated with S-2P, followed by a fixed concentration of the competitor. Levels of bound competitors were measured as the final readout. (**D**) Per-mouse serum competition from week 8 against hDPP4-Fc binding to S-2P. Background serum reactivity to *top,* secondary antibody; *middle,* competition between hDPP4-Fc and serum; and *bottom,* normalized data obtained by subtracting background secondary antibody signal from competition. The arithmetic mean of the normalized data points at each dilution are shown in Fig. 3A. (**E**) Per-mouse serum competition from week 8 against G2 binding to S-2P, processed as in (D).

**Fig. S5.**
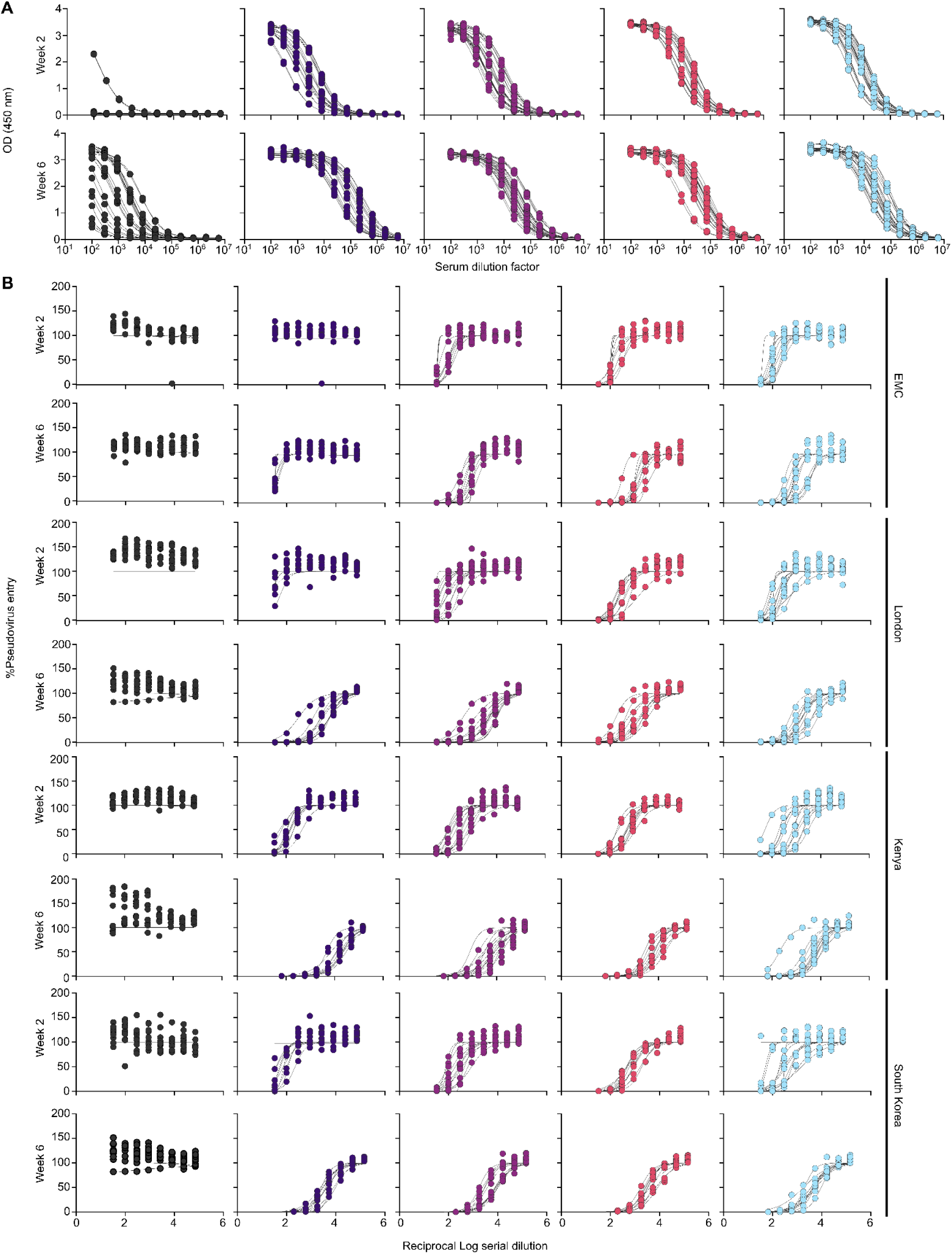
288/330^+/+^ mouse serum binding to MERS-CoV S-2P and pseudovirus neutralization. (**A**) Raw week 2 and 6 serum ELISA data against vaccine-matched (EMC) MERS-CoV S-2P. (**B**) Raw neutralization data from weeks 2 and 6 using pseudoviruses bearing the MERS-CoV EMC, London, Kenya, and South Korea variant spikes.

**Table 1.**
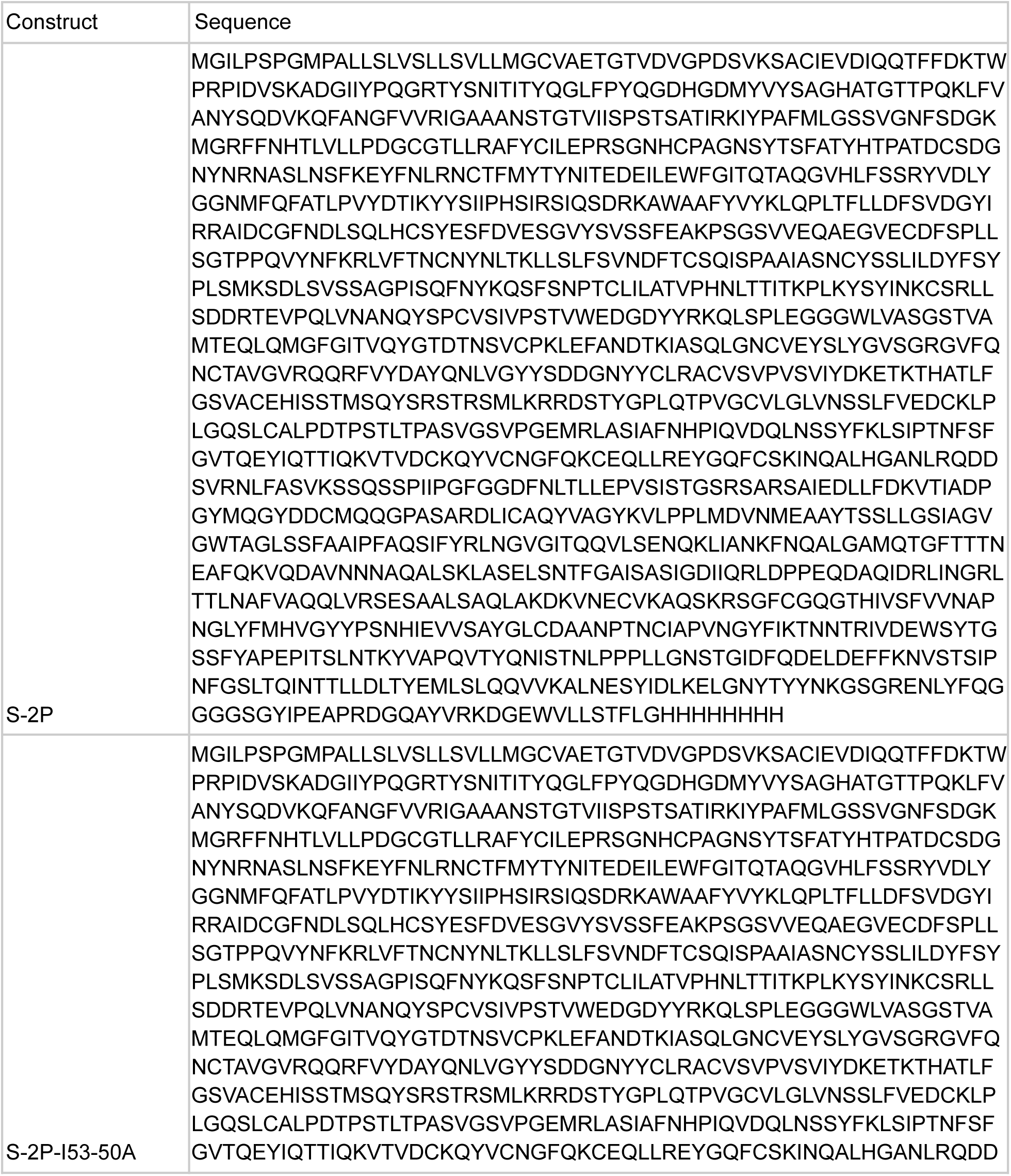
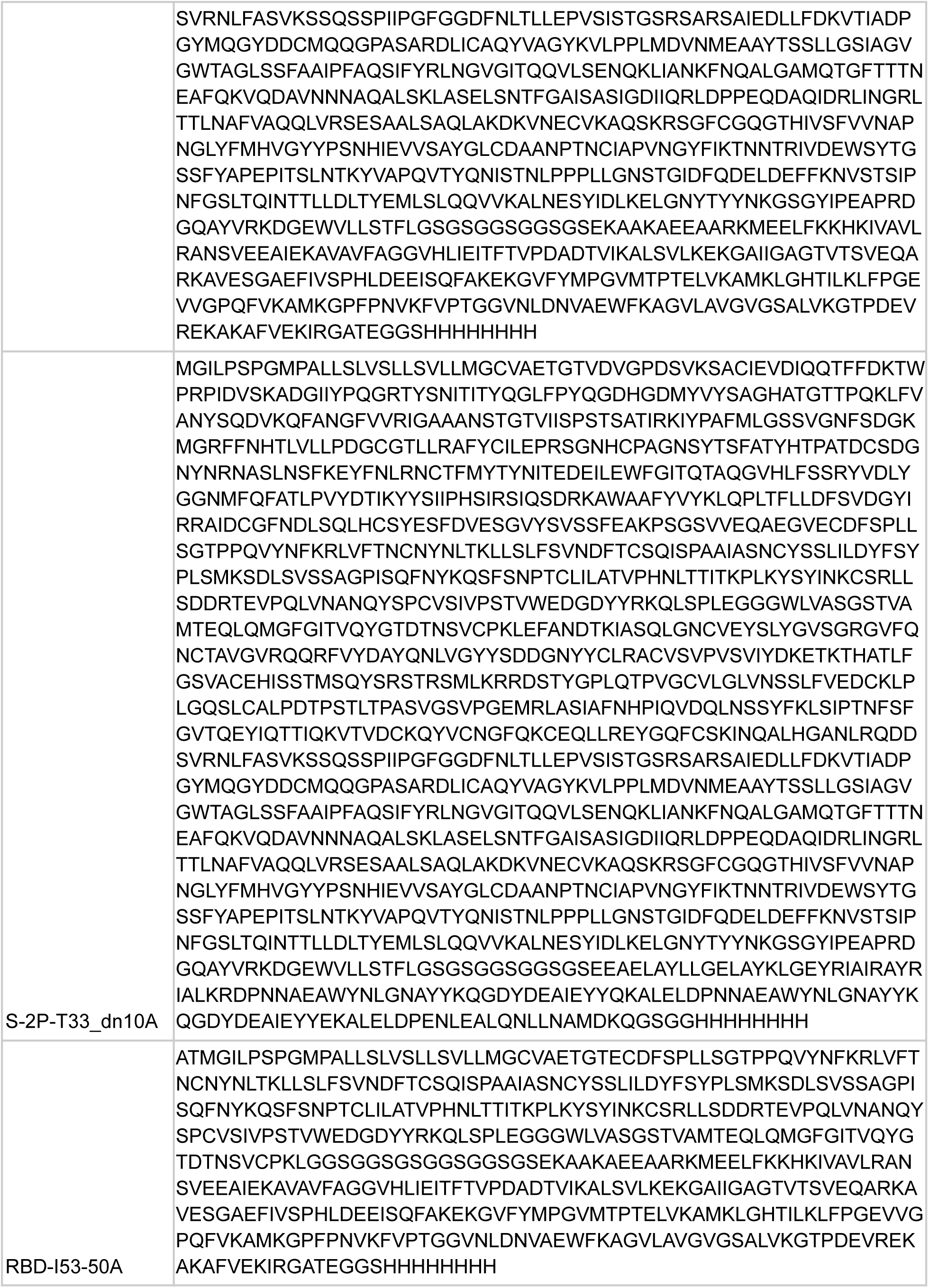
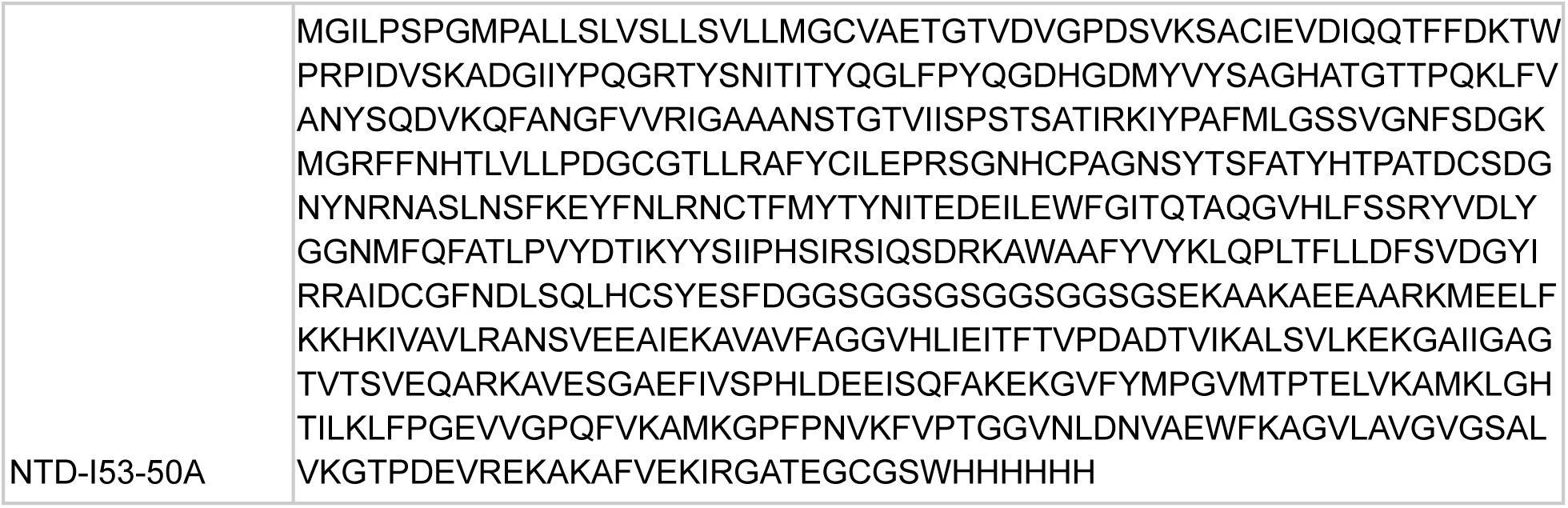
Amino acid sequences of novel proteins used in this study.

